# Defining compartmentalized stem and progenitor populations with distinct cell division frequency in the ocular surface epithelium

**DOI:** 10.1101/2020.06.23.156505

**Authors:** Ryutaro Ishii, Hiromi Yanagisawa, Aiko Sada

## Abstract

Adult tissues contain label-retaining cell (LRC)s, which are relatively slow-cycling and considered to represent a unique property of tissue stem cell (SC)s. In the ocular surface epithelium, LRCs are detected in the limbus, a boundary between cornea and conjunctiva, and the fornix of the conjunctiva; however, the character of LRCs and identity of SCs remain unclear due to lack of appropriate molecular markers. Here we show that the ocular surface epithelium accommodates spatially distinct stem/progenitor populations with different cell division frequency. By combining EdU pulse-chase analysis and lineage tracing with three CreER transgenic mouse lines: Slc1a3^CreER^, Dlx1^CreER^ and K14^CreER^, we detect distinct dynamics of epithelial SCs in the cornea and conjunctiva. In the limbus, long-lived SCs are labeled with Slc1a3^CreER^ and they either migrate centripetally toward the central cornea or laterally expand their clones within the limbal region. In the central cornea, cells are mostly non-LRCs, labeled by Dlx1^CreER^ and K14^CreER^, and the number of clones declines after a short period of time with rare long-lasting clones, suggesting their properties as short-lived progenitor cells. In the conjunctival epithelium, which consists of bulbar, fornix and palpebral conjunctiva, each territory is regenerated by compartmentalized, distinct SC populations without migrating one region to another. The severe damage of the cornea leads to the cancellation of SC compartments, causing conjunctivalization of the eye, whereas milder limbal injury induces a rapid increase of laterally-expanding clones in the limbus. Taken together, our work provides lineage tracing tools of the eye and defines compartmentalized, multiple SC/progenitor populations in homeostasis and their behavioral changes in response to injury.

## Introduction

Tissue stem cell (SC)s have the ability to self-renew and differentiate and play an important role in homeostasis and injury repair. Adult epithelial tissues, such as the skin, eye, oral mucosa and intestine, show proliferative heterogeneity. In hair follicles, for example, infrequently-dividing or “slow-cycling” cells in the bulge region are identified as label-retaining cell (LRC)s by pulse-chase experiments and show unique, long-lived stem cell property (Tumbar, et al., 2004; Cotsarelis, et al., 1990; Bickenbach, 1981). The hierarchical stem/progenitor model, in which slow-cycling SCs give rise to short-lived, fast-dividing progenitor or so-called transit-amplifying cells, has been applied to various epithelial or non-epithelial tissues (Sanchez-Danes, et al., 2016; Mascre, et al., 2012; Foudi, et al., 2009; Fuchs, 2009; Sangiorgi and Capecchi, 2008; Wilson, et al., 2008). However, studies have challenged the generality of the stem/progenitor model and suggested that a relationship between LRCs and their SC potential can be tissue-or context-dependent. In the intestine, a fast-dividing Lgr5^+^ population is suggested to act as SCs (Barker, et al., 2007)and LRCs represent committed precursor cells (Buczacki, et al., 2013). In the interfollicular epidermis (IFE) and oral epithelium, different epithelial compartments accommodate heterogeneous populations of SCs, which show differences in cell division frequency, location, molecular property, biological functions or tumorigenic abilities (Byrd, et al., 2019; Kretzschmar, et al., 2016; Roy, et al., 2016; Sada, et al., 2016; Sanchez-Danes, et al., 2016; Fullgrabe, et al., 2015; Gomez, et al., 2013; Page, et al., 2013). On the other hand, a single population model suggested that epithelial tissues are maintained by an equipotent, single-cell population, which undergoes stochastic divisions and fate choices, not by the existence of functionally discrete cell populations (Jones, et al., 2019; Krieger and Simons, 2015; Doupe, et al., 2012; Rompolas, et al., 2012; Doupe, et al., 2010; Clayton, et al., 2007)

The ocular surface epithelium consists of the cornea and the conjunctiva and protects the eye from environmental damages. The cornea is covered by stratified, non-keratinizing squamous epithelium, which lies on the avascular corneal stroma. The conjunctival epithelium is comprised of three parts (bulbar, fornix, and palpebral conjunctiva) and works as a physical protective barrier as well as secretes mucin required for a tear film homeostasis through goblets cells (Hertsenberg and Funderburgh, 2015; Lavker, et al., 2004). Severe corneal injury and loss of stem cells lead to an invasion of conjunctival cells to the cornea (conjunctivalization), resulting in the corneal opacity associated with neovascularization and eventually visual loss.

The pulse-chase experiments using histone H2B-GFP, BrdU or tritiated thymidine have suggested the existence of LRCs in the limbus and fornix regions of the conjunctiva (Parfitt, et al., 2015; Wei, et al., 1995; Cotsarelis, et al., 1989b). The limbus was proposed as the region containing a unique SC population, known as limbal epithelial SCs that give rise to progenitors and migrate toward the central cornea (Cotsarelis, et al., 1989a) (Lavker and Sun, 2003). Limbal epithelial SCs showed the holoclone-forming ability in vitro (Pellegrini, et al., 1999; Ebato, et al., 1988) and used as a source for regenerative therapy of the corneal epithelium (Rama, et al., 2010). The limbal SC model is supported by lineage tracing studies using inducible Cre-mediated labeling. K14^CreER^ marked cells were observed migrating centripetally from the limbus to the central cornea (Richardson, et al., 2017; Richardson, et al., 2016; Amitai-Lange, et al., 2015; Di Girolamo, et al., 2015). The CAGG^CreER^, which labels all cell types in the corneal epithelium, showed long-lived clones emerged from the limbus, but not in the central cornea (Dora, et al., 2015). Bmi1^CreER+^ has been shown to label relatively shorter-lived progenitor populations located in the central cornea (Kalha, et al., 2018).

The alternative model suggested the existence of SCs in the central cornea, known as corneal epithelial SC hypothesis (Majo, et al., 2008). In this model, corneal epithelial SCs that were transplanted to the limbus migrated to central cornea when the entire cornea was removed. In addition, the corneal epithelial cells exhibited the ability to undergo serial transplantation, suggesting that self-renewing SCs exist in the entire corneal epithelium. The analysis of K14^CreER^-based lineage tracing, transplantation and culture experiments also support the existence of SC/progenitor population in the central or entire cornea (Li, et al., 2017; Amitai-Lange, et al., 2015). However, lack of lineage tracing tools to specifically mark epithelial subpopulations and definitive regional markers hampered us to determine the SC identity of the corneal epithelium.

Despite accumulating knowledge of corneal regeneration, the character of conjunctival SCs has been less explored (Ramos, et al., 2015). A theory of conjunctival transdifferentiation proposed that conjunctival epithelial cells may migrate and become corneal epithelium (Shapiro, et al., 1981). In contrast, studies have shown that conjunctival and corneal epithelial cells exhibit distinct intrinsic property and differentiation potential in the same environmental conditions, ruling out the possibility of conjunctival transdifferentiation (Cho, et al., 1999; Wei, et al., 1996; Wei, et al., 1993). Based on the location of LRCs and in vitro holoclone-forming ability, fornix conjunctiva has been proposed to contain conjunctival epithelial SCs (Wei, et al., 1995; Wei, et al., 1993). Other studies suggested the bulbar conjunctiva (Budak, et al., 2005; Pellegrini, et al., 1999) or palpebral conjunctiva (Chen, et al., 2003) as an epithelial SC location. Due to a lack of genetic mouse tools, in vivo evidence is missing as to which populations of conjunctival epithelium act as SCs and what is the lineage relationship among three regions of the conjunctiva (bulbar, fornix and palpebra).

Studies in skin and other epithelial tissues have shown that epithelial SCs display plasticity to respond to tissue damages and change their lineages, transiently or permanently (Dekoninck and Blanpain, 2019; Belokhvostova, et al., 2018). In eyes, Nasser et al. combined K14^CreER^ with K15^GFP^ reporter and proposed that the limbus epithelium deletion is repaired by dedifferentiation of corneal committed cells (Nasser, et al., 2018). In contrast, after chemical burn or whole cornea epithelium deletion, the cornea is covered with conjunctiva-like epithelium in accordance with vascularization in the stroma (Afsharkhamseh, et al., 2016; Saika, et al., 2005; Wei, et al., 1995). It remains to be addressed how different subpopulations of SCs in the cornea (central vs peripheral) and conjunctiva (bulbar vs fornix vs palpebra) react to different levels of tissue damage.

The previous studies showed the limbus is molecularly defined by their high level of p63, K15 and Abcb5 (Sartaj, et al., 2017; Pellegrini, et al., 2001). Recent RNA seq studies provided the whole transcriptome of H2B-GFP LRCs (Sartaj, et al., 2017) or the entire ocular surface epithelium by single cell analysis (Kaplan, et al., 2019). However, no definitive markers for lineage tracing have been found to faithfully label and distinguish the limbus and other populations of corneal epithelium. In our previous study, we identified two markers, Dlx1 and Slc1a3, which preferentially label LRC SCs and non-LRC SCs in the IFE of skin, respectively (Sada, et al., 2016). These two populations of IFE SCs are largely independent of each other during homeostasis, replenishing their own compartments, but they also show plasticity to contribute to each other’s lineage in response to injury. It remains unknown whether such SC compartments also exist in the cornea or conjunctiva and can be marked by LRC and non-LRC SC markers established in other epithelial tissues. Here, we applied lineage tracing tools, Dlx1^CreER^, Slc1a3^CreER^, and K14 ^CreER^ to ocular surface epithelium and characterized cellular dynamics in homeostasis and injury. We showed that each Cre labeled distinct populations of stem and progenitor cells that were highly compartmentalized and different in cell division frequency. Under physical or chemical damage, these territorial segregations were lost and SC lineages were altered. These findings give a new insight into the understanding of biological natures of ocular epithelial SCs.

## Results

### LRCs and non-LRCs in the ocular surface epithelium are marked by distinct CreER tools, K14 ^CreER^, Dlx1^CreER^ and Slc1a3^CreER^

Previously, LRCs were shown to localize in the limbus and fornix conjunctiva (Parfitt, et al., 2015). To evaluate the distribution of LRCs in the whole eye, we re-analyzed LRC locations in detail by applying whole-mount techniques of epithelial sheets of the skin IFE (Braun, et al., 2003) (Figure 1A, B). A nucleotide analogue EdU is incorporated into all dividing cells, whether or not they are stem cells, during 1 week of treatment (=pulse) (Figure 1C). Since the label is lost by divisions or differentiation during 5 weeks of chase, only cells that divide infrequently can be detected as LRCs. The corneal and conjunctival epithelium were demarcated by K12 and K19, respectively (Figure S1A) (Braun, et al., 2003). At 0-day-chase, EdU+ cells were entirely distributed in the cornea and conjunctiva (Figure. 1E). At 5-week-chase, EdU+ cells were preferentially enriched in the limbus, which was at the boundary of K12-positive and -negative area, and the fornix area in the center of conjunctiva. This result confirmed the distribution pattern of LRCs in the ocular surface epithelium.

**Figure 1.**
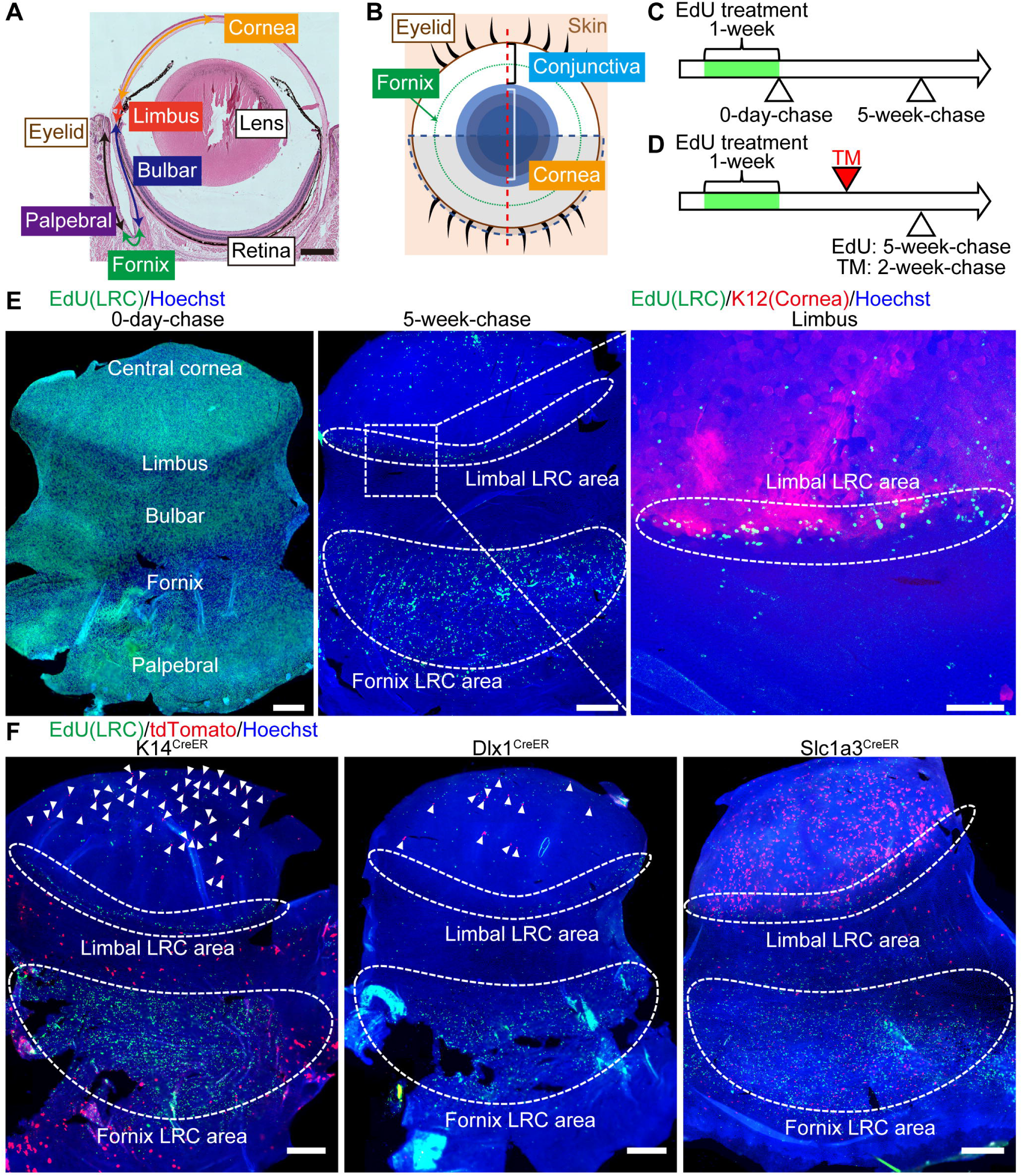
Distinct CreER tools mark label-retaining cell (LRC) and non-LRC compartments. (A) Hematoxylin-eosin stained mouse eye. Scale bar: 500 μm. (B) A schematic representation of the mouse eye. The eye is analyzed by sagittal sections (red dotted line) or whole-mount preparation of the epithelium (blue dotted line). (C) EdU pulse-chase scheme to detect LRCs in the ocular surface epithelium. (D) Scheme to examine the relationship of CreER-marked cells with LRCs and non-LRCs. TM, Tamoxifen. (E) A whole-mount staining of epithelial sheets after EdU pulse-chase experiments at 0-day-(left) and 5-week-(center) chases. Limbal areas are shown with higher magnification (right). The dashed line surrounds LRC area. Green, EdU. Red, K12 (corneal marker). Blue, Hoechst. Scale bars: 500 μm (E, left and center), 200 μm (E, right). (F) A whole-mount staining of epithelial sheets after Tamoxifen injection and EdU pulse-chase in K14^CreER^, Dlx1^CreER^ and Slc1a3^CreER^. The dashed line surrounds LRC area. Arrowheads indicate tdTomato+ cells in the cornea. Green, EdU. Red, tdTomato. Blue, Hoechst. Scale bars: 500 μm.

Next, we analyzed the relationship between LRC distribution and three CreER: K14^CreER^, Dlx1^CreER^ and Slc1a3^CreER^. According to previous reports using K14^CreER^-Confetti mice, K14^CreER^ labeled cells were uniformly distributed in the cornea and conjunctiva (Lobo, et al., 2016; Di Girolamo, et al., 2015). In our study, we used a different strain of K14^CreER^ with relatively lower K14 promoter activity, and injected a reduced dose of Tamoxifen to detect subpopulations of epithelial cells as was performed in skin (Zhang, et al., 2010). In addition to K14^CreER^, we used Dlx1^CreER^ and Slc1a3^CreER^, which we previously established as IFE SC markers: Dlx1 marks LRCs and Slc1a3 marks non-LRCs (Sada, et al., 2016). We used the same EdU pulse-chase condition to detect LRCs and injected Tamoxifen at 2 weeks before analysis (Figure 1D and Figure S1B-D). No tdTomato expression was observed without Tamoxifen injection in any CreER used (Figure S1E). We found that Slc1a3^CreER^ labeled cells in the limbal LRC region as well as peripheral cornea, whereas K14^CreER^ and Dlx1^CreER^ preferentially labeled the central cornea (Figure 1F). In conjunctiva, LRC-dense fornix region was preferentially marked by Slc1a3^CreER^ and the bulbar and palpebral conjunctiva were marked by K14^CreER^. These results suggest the possible heterogeneity of ocular surface epithelium in their cell division frequency and molecular characters. K14^CreER^, Dlx1^CreER^ and Slc1a3^CreER^, can be used as genetic tools to distinguish cells within LRC and non-LRC regions in the cornea and conjunctiva.

### Slc1a3^CreER^ marks limbal SC populations with two distinct dynamics

To analyze the behavior of LRC population in the limbus, we used Slc1a3^CreER^ for lineage tracing. At 2-week-chase, tdTomato-labeled cells were predominantly observed in the basal layer of limbus and peripheral cornea (Figure 2A-C). To quantitatively analyze the distribution of labeled clones, we measured the length between the corneal/conjunctival boundary to proximal edge of each clone and plotted in the histogram (Figure S1F). The boundary was determined by K12 or K19 staining of whole-mount images. We found that the location of Slc1a3^CreER+^ clones at 2-week-chase peaked within 0-500 μm, with a gradual decline toward the central cornea (Figure 2D). By 1-month-chase, the labeled cells started to show radial stripes, indicating the continuous migration and expansion of cells from the limbus toward the central cornea as previously reported (Richardson, et al., 2017; Richardson, et al., 2016; Amitai-Lange, et al., 2015; Di Girolamo, et al., 2015; Dora, et al., 2015). The clones reached to the upper most layers of epithelium after 1-month-chase, showing their differentiation ability (Figure 2A-D). At 3-month-chase, the distribution of clones shifted toward the central cornea and peaked ∼750-1000 um from the boundary (Figure, 2D), indicating some clones were short-lived and lost within a few months. Notably, the labeled cells at the limbal region showed at least two distinct behaviors; (1) the radial stripes extended from the limbus (Figure 2B, white arrowheads, radial stripe type), and (2) the clones expanded within the limbal region (Figure 2B, yellow arrowheads, laterally-expansion type). The former was located at K12-positive corneal area, and the latter was located at the boundary or K12-negative area. At 1-year-chase, ∼80% of clones, both radial stripe type and laterally-expansion type, were maintained as independent clones (Figure, 2A-D and Figure S2). The laterally-expansion type of clones were relatively small in size and became apparent after long-term chase, indicating their slow-cycling, infrequently-dividing nature. Taken together, Slc1a3^CreER^ lineage tracing proposed that limbal LRC region contains long-lived SCs that undergo either centripetal migration to replenish corneal epithelium or lateral expansion to maintain limbal compartment with relatively slow turnover.

**Figure 2.**
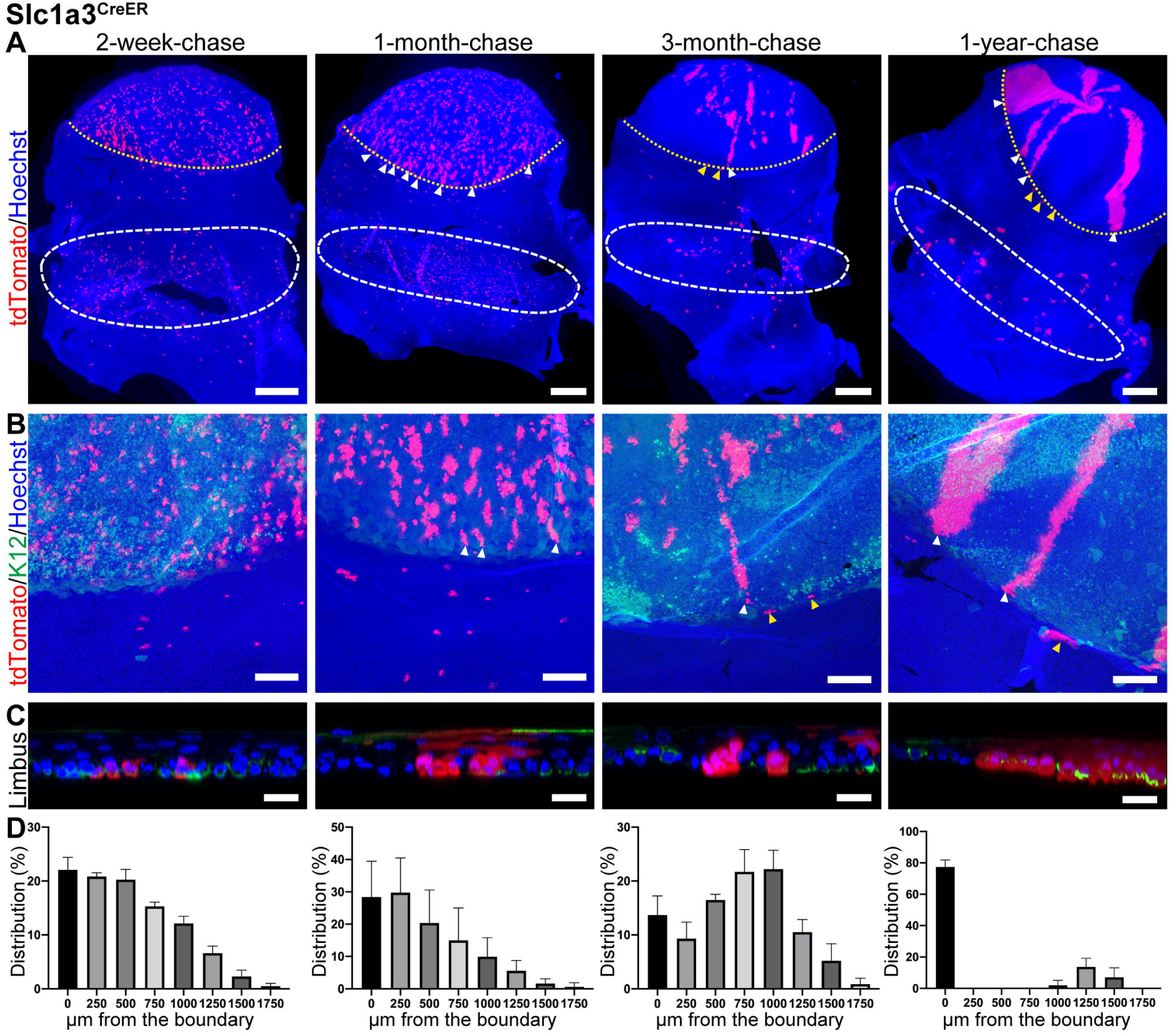
Lineage tracing of Slc1a3^CreER^ in the limbus and cornea. (A-C) Whole-mount immunostaining at 2-week-, 1-month-, 3-month- and 1-year-chases. The yellow dashed line represents corneal/conjunctival boundary (A). The white dashed line represents tdTomato+ cell-enriched area in the fornix conjunctiva (A). Limbal areas are shown with higher magnification (B) and side view of Z-stack confocal image (C). White arrowheads indicate tdTomato+ radial stripes extended from the limbus (A, B). Yellow arrowheads indicate tdTomato+ clones expanded laterally within the limbal region (A, B). Red, tdTomato. Green, K12 (B, C, corneal marker). Blue, Hoechst. Scale bars: 500 μm (A), 200 μm (B), 20 μm (C). (D) Distribution of the length of tdTomato+ clones from the corneal/conjunctival boundary expressed as the percentage of total clones. *N* ≥ 3. All tdTomato+ clones in whole-mount samples from a half eye are measured and used for quantification. Error bars show S.E.M.

### K14^CreER +^ and Dlx1^CreER+^ short-lived progenitor populations in the central cornea

To address whether cells in the central cornea retain the SC ability in vivo, we traced the fate of K14^CreER^ and Dlx1^CreER^ marked cells, which were preferentially observed in non-LRC, central corneal region (Figure 1F). With this tool, we can determine whether labeled cells are able to self-maintain themselves for long periods in accordance with the corneal epithelial SC hypothesis, or are supplied from limbal epithelial SCs. At 2-week-chase, K14^CreER^ marked clones preferentially located in the basal layer of the central cornea (Figures 3A-C). A quantitative analysis showed that the peak of histogram was ∼1000 µm away from the boundary (Figure 3D), which is distinct from the distribution of Slc1a3^CreER+^ clones (Figure 2D). During 3 months of chase, the clone distribution was slightly shifted toward the central cornea and peaked at ∼1500 µm. These observed clones in the central cornea remained until 3 months of chase and the number of clones were markedly decreased by 1 year of chase (Figure 3E). At 1-year-chase, remaining clones in the central cornea consisted of basal and a few suprabasal cells, which may reflect their limited ability to differentiate (Figures 3A-C). Although the labeling efficiency is much lower than K14^CreER+^, the lineage tracing by Dlx1^CreER^ showed similar labeling patterns and cellular dynamics (Figure S3). These results suggested that the central cornea, the non-LRC territory of the cornea, contained shorter-lived progenitor populations, which were marked by K14^CreER^ and Dlx1^CreER+^.

**Figure 3.**
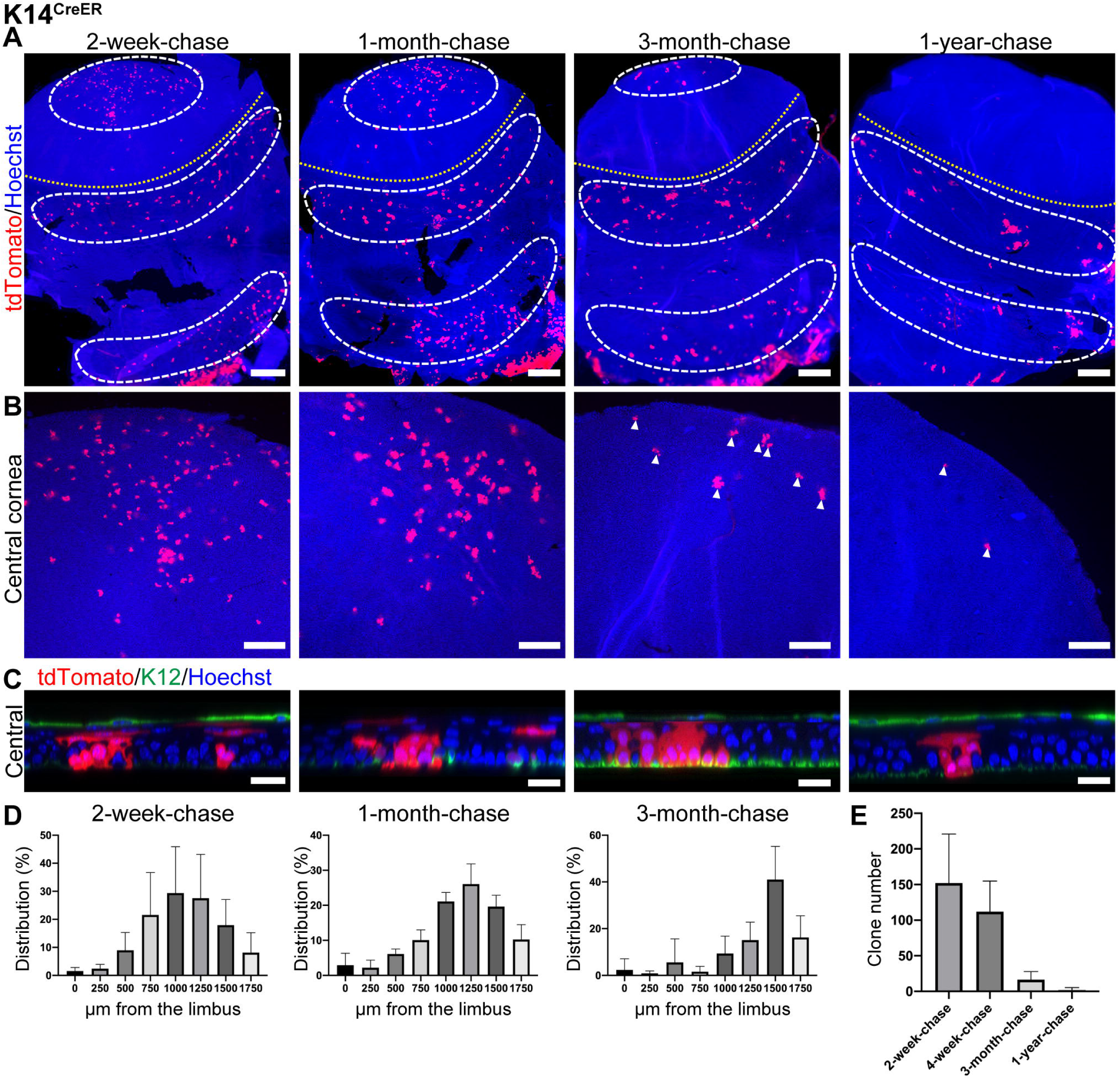
Lineage tracing of K14^CreER^ in the cornea. (A-C) Whole-mount immunostaining at 2-week-, 1-month-, 3-month- and 1-year-chases. The yellow dashed line represents corneal/conjunctival boundary (A). The white dashed line represents tdTomato+ cell-enriched area (A). Central corneal areas are shown with higher magnification (B) and side view of Z-stack confocal image (C). Arrowheads indicate tdTomato+ cells (B). Red, tdTomato. Green, K12 (B, C, corneal marker). Blue, Hoechst. Scale bars: 500 μm (A), 200 μm (B), 20 μm (C). (D) Distribution of the length of tdTomato+ clones from the corneal/conjunctival boundary expressed as the percentage of total clones. (E) Number of tdTomato+ clones per a half whole mount sample is quantified at indicated time points. *N* ≥ 3. All tdTomato+ clones in whole-mount samples from a half eye are measured and used for quantification. Error bars show S.E.M.

### Three distinct SC populations in the conjunctiva maintained their own compartments

The SC identity in the conjunctiva remains elusive. To define SC location and their cellular lineages in the conjunctiva, we used Slc1a3^CreER^ as a marker of LRC region in the fornix, and K14^CreER^ as a marker of non-LRC region in the bulbar and palpebral conjunctiva (Figure 1F). We first tested whether Slc1a3^CreER+^ LRC population in the fornix acts as SCs and which epithelial compartments are maintained by this population. We found that the clones marked by Slc1a3^CreER^ in the fornix conjunctiva were located at ∼1500 μm away from the corneal/conjunctival boundary toward the eyelid and consisted of a small cluster of basal cells at 2-week-chase (Figure 2A and Figure 4A, C). These clones expanded in size and remained in the same region during 1 year of chase (Figure 2A and Figure 4A-C). No apparent migration was found from the fornix conjunctiva to other regions, including bulbar, palpebral conjunctiva or cornea, indicating that fornix LRCs are long-lived SCs that regenerate their own compartment. To further address which cells contribute to the regeneration of bulbar and palpebral conjunctival epithelium, we preformed lineage tracing using K14^CreER^, which preferentially labels non-LRCs in the bulbar and palpebral conjunctiva (Figure 3A). K14^CreER+^ showed enrichment of clones peaked at ∼250 μm (bulbar conjunctiva) and ∼2250 μm (palpebral conjunctiva) away from the boundary, which are distinct from that of Slc1a3^CreER^. After chase, K14^CreER+^ clones in the bulbar or palpebral conjunctiva showed no directed movement and expanded at their own territories (Figure 4D-F). Thus, these results suggest that three distinct SC populations were located in the conjunctiva and maintained in their own compartments, excluding the possibilities that fornix LRCs are the sole source of SC population to reconstitute the entire conjunctiva or transdifferentiate into the corneal epithelium.

**Figure 4.**
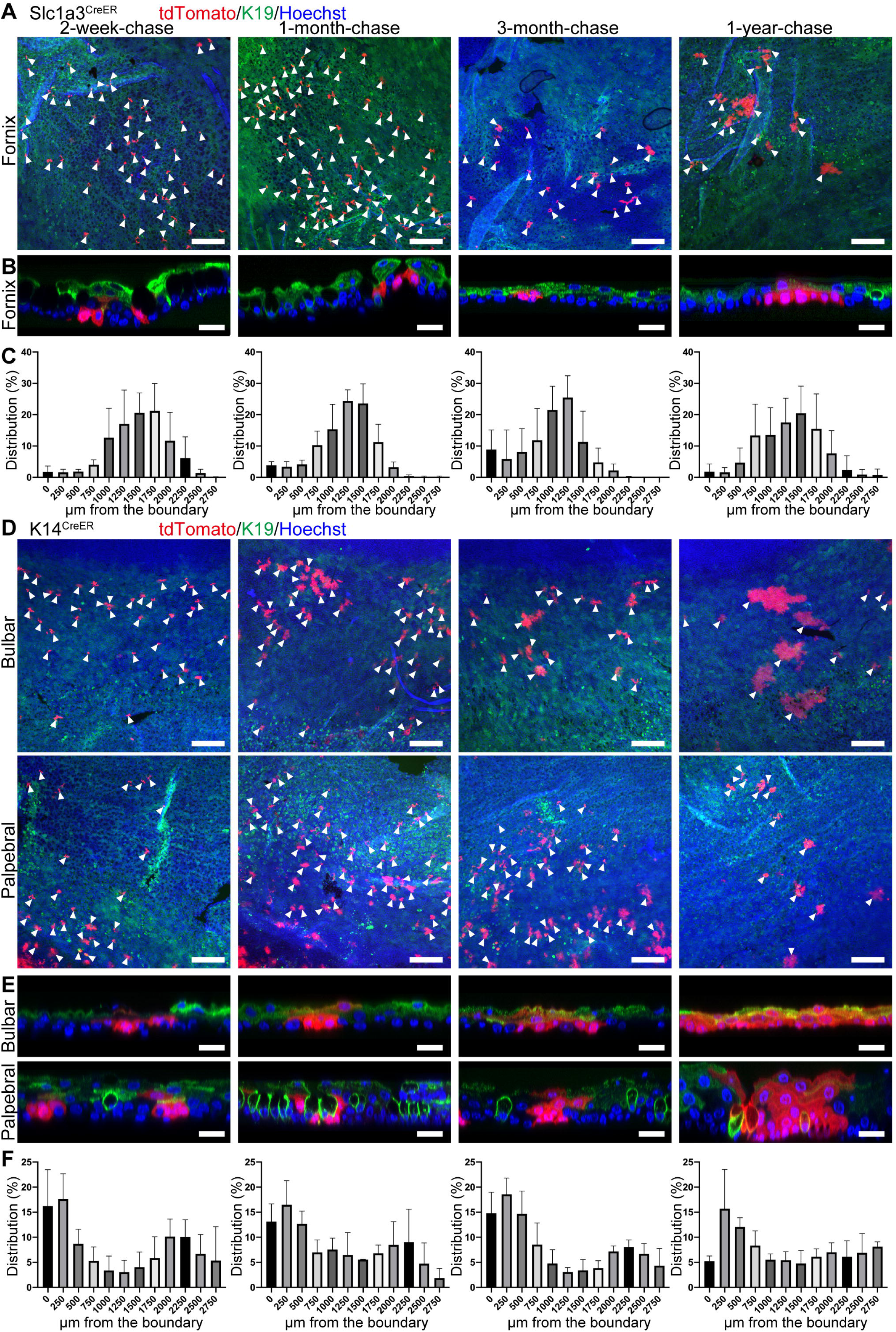
Lineage tracing of Slc1a3^CreER^ and K14^CreER^ in the conjunctiva. (A, B, D, E) Mouse eyes are analyzed by whole-mount immunostaining at 2-week-, 1-month-, 3-month- and 1-year-chases. Fornix, bulbar and palpebral conjunctiva are shown as projected image (A, D) or side view of Z-stack confocal image (B, E). Arrowheads indicate tdTomato+ cells (A, D). Red, tdTomato. Green, K19 (conjunctival marker). Blue, Hoechst. Scale bars: 200 μm (A, D), 20 μm (B, E). (C, F) Distribution pattern of tdTomato+ clones. Length from the corneal/conjunctival boundary to the center of each clone are measured. *N* ≥ 3. All tdTomato+ clones in whole-mount samples from a half eye are measured and used for quantification. Error bars show S.E.M.

Although the overall distribution of Slc1a3^CreER+^ and K14^CreER+^ clones remained constant over time (Figure 4C, F), we found that the number of K14^CreER+^ bulbar conjunctival clones nearest to the boundary (0∼250 μm) decreased at 1-year-chase (Figure 4F). These clones may be replaced by the laterally-expanded Slc1a3^CreER+^ LRC population in the distinct limbal compartment (Figure 2B, yellow arrowheads), which most likely represent long-lived, slow-cycling SCs.

### Injury triggers remodeling of SC compartments in the ocular surface epithelium

It has been shown in skin and eye that epithelial SCs have the potential to alternate their behavior in a context dependent manner. To test how different SC/progenitor populations in the cornea and conjunctiva respond to the injury, we applied two types of injury: limbal epithelial deletion and chemical burn, and analyzed the behavior of cells residing in the distinct epithelial compartments. Previous reports suggested that limbal SC deletion is recovered by de-differentiation and migration of progenitor cells located in the cornea (Nasser, et al., 2018). We took advantage of Slc1a3^CreER^ and K14^CreER^ to distinguish cells in the limbus, peripheral, central cornea and different compartments of conjunctiva and to see which populations are the source of limbal regeneration. At two weeks after Tamoxifen injection, we performed limbal epithelial injury by deleting the epithelium located above the vessels using an ophthalmic rotating burr (Figure 5A). The limbal epithelial deletion was confirmed by fluorescein staining (Figure 5B). Before limbal deletion, Slc1a3^CreER^ marked cells were located in the limbus and peripheral cornea (Figure 5C). In contrast, K14^CreER^ marked cells were located in the central cornea and the bulbar conjunctiva (Figure 5D). At 1-week post injury, Slc1a3^CreER^ marked clones were started to expand laterally at K19-positive region (Figure 5C and Figure S4A, yellow arrowheads). This population possibly corresponds to the laterally-expanding clones that we observed at 3 months and 1 year of chase during normal homeostasis (Figure 2B and Figure S2, yellow arrowheads), and these clones appeared quickly after the injury. At 4-week-post-injury, the radial stripes were reappeared at the K19-negative corneal region (Figure S4A). In contrast, K14^CreER^ marked cells, either at the central cornea or the bulbar conjunctiva, didn’t show such behavior change upon injury (Figure 5D and Figure S4A). Thus, it appears that slow-cycling Slc1a3^CreER+^ cells in the K19-positive limbal compartment in the vicinity of injury were rapidly activated after injury and contributed to the recovery of the limbal epithelium. In contrast, central cornea or bulbar conjunctiva marked by K14^CreER^ did not participate in the repair process. Our results suggest behavior differences between Slc1a3^CreER+^ (limbus/peripheral cornea) vs K14^CreER+^ (central cornea/bulbar conjunctiva) cell populations in response to limbal epithelial injury.

**Figure 5.**
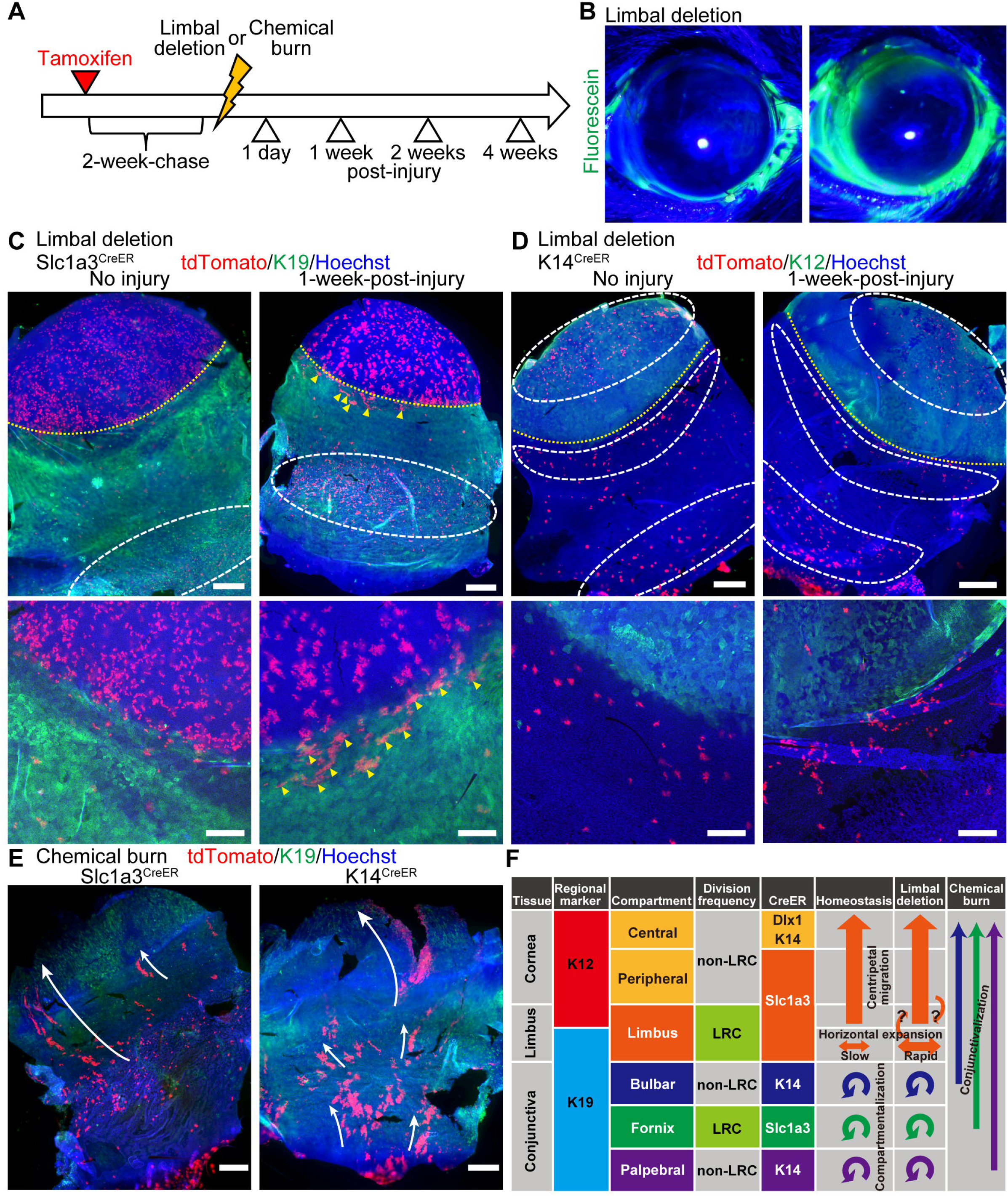
Dynamics of Slc1a3^CreER^ and K14^CreER^ population after injury. (A) Experimental scheme of injury experiments. (B) Fluorescein staining of eye at pre-injury (left) and post-injury (right). (C, D) Whole-mount immunostaining after limbal epithelial deletion. Control (left) or 1-week-post-injury (right) are shown. The yellow dashed line represents corneal/conjunctival boundary. The white dashed line represents dTomato+ cell-enriched area. Yellow arrowheads indicate tdTomato+ clones expanded laterally within the K19-positive region (C). Red, tdTomato. Green, K19 (C, conjunctival marker), K12 (D, corneal marker). Blue, Hoechst. Scale bars: 500 μm (top), 200 μm (bottom). (E) Whole-mount immunostaining at 1-week post chemical burn. Slc1a3^CreER^ (left) or K14^CreER^ (right) marked cells are shown. White arrow represents movement of conjunctival tdTomato+ clones. Red, tdTomato. Green, K19 (conjunctival marker). Blue, Hoechst. Scale bars: 500 μm. (F) Proposed model of compartmentalized stem and progenitor populations with distinct cell division frequency in the ocular surface epithelium.

Finally, we introduced chemical injury by applying sodium hydroxide solution to the cornea (Saika, et al., 2005). Stromal injury by alkali burn leads to limbal SC deficiency, including conjunctivalization of the corneal surface and neovascularization (Joussen, et al., 2003). However, direct evidence is lacking as to which conjunctival population responds to the corneal damage. After alkaline burn, the cornea was largely covered by the conjunctival cells, which expressed conjunctival marker K19 in the entire epithelium and lost K12 expression (Figure 5E and Figure S4B). By using Slc1a3^CreER^, we found that epithelial SCs in the fornix conjunctiva migrated out from their territory and expressed K19 in the corneal area. This lineage disruption persisted for 4 weeks (Figure S4B). Similarly, K14^CreER^ marked clones, both in the bulbar and palpebral conjunctiva, migrated toward the cornea (Figure 5E and Figure S4B). These observations indicate that multiple conjunctival SCs lost their identity and territorial distribution pattern after the extensive mesenchymal damage and migrated out of their territories to compensate for the loss of limbal SCs.

## Discussion

Slow-cycling nature has been linked to SC property; however, the character of these cells in the ocular surface epithelium has been controversial due to lack of definitive molecular markers and lineage tracing mouse models. In our current study, taking advantage of three CreER tools, we analyzed the cellular dynamics of corneal and conjunctival epithelium during homeostasis and injury repair (Figure 5F). We define distinct compartments in the ocular surface epithelium, which are characterized by the marker expression, cell division frequency and anatomical location. Slc1a3^CreER^ marker preferentially labels LRC regions in the limbus and fornix conjunctiva, which are distinct from labeling pattern of K14^CreER^ or Dlx1^CreER^ lines. Notably, three compartments in the conjunctiva, bulbar, fornix and palpebral conjunctiva, are governed by its local SC populations marked by distinct CreER tools, indicating their differences in molecular property. Chemical burn triggers disruption of SC compartments and invasion of all three conjunctival SC populations into the corneal region. The territorial segregation of epithelial SCs is determined by yet unidentified mechanism and possibly involves stromal architecture, extracellular matrix and secreted factors. Our work provides genetic tools to precisely mark and examine the dynamic behavior of multiple SC/progenitor populations in the ocular surface epithelium during homeostasis and injury repair, and to molecularly characterize each population.

In the ocular surface epithelium, we found Slc1a3^CreER^ activity is enriched in LRC regions but its functional importance remains to be addressed. Slc1a3 gene encodes a glutamate/aspartate transporter and functions for glutamatergic neurotransmission in brain (Kanai and Hediger, 2004). In peripheral tissues, Slc1a3 is widely expressed in epithelial cells (Berger and Hediger, 2006). Slc1a3 is up-regulated in actively-cycling SCs of skin IFE, hair follicles and sebaceous gland and plays a role in SC/progenitor cell activation (Reichenbach, et al., 2018; Sada, et al., 2016). Slc1a3 promotes cell proliferation and survival of cancer cells under nutrient starvation or hypoxic environment (Garcia-Bermudez, et al., 2018; Tajan, et al., 2018). Slc1a3 also mediates tumor growth by exchanging glutamate and aspartate between squamous cell carcinoma and carcinoma-associated fibroblasts in stiff environment (Bertero, et al., 2019). Given that proliferative heterogeneity of SCs is highly correlated with metabolic regulation (Coller, 2019; Coloff, et al., 2016), it will be interesting to address the roles of Slc1a3 and amino acid metabolism in SCs and their niches of ocular surface epithelium.

Slc1a3^CreER+^ SC population in the limbus shows two distinct behavioral patterns: one migrated centripetally toward the central cornea; the other self-expanded laterally within the limbal compartment. Our analysis cannot fully address whether these two types of Slc1a3^CreER+^ clones represent two discrete SC populations or differences in cell division pattern (asymmetric vs symmetric) of the same SC population. During steady-state tissue turnover, the dynamics of the limbal SC is biased toward the production of corneal epithelial progenitors in the central cornea. The limbal injury, however, shifts dynamics toward the limbal-expansion mode and induces rapid expansion of the Slc1a3^CreER+^ population within the limbus. Hence, Slc1a3^CreER+^ population in the limbus, despite relatively slow-cycling in nature, equip itself with a back-up system to quickly respond to tissue damage. Our data does not rule out the possibility that de-differentiation of peripheral corneal progenitor cells may contribute to the limbal SC regeneration. Further studies to track Slc1a3^CreER+^population by different methods, e.g., live imaging during the process of injury repair, are needed to address this point.

The K14^CreER^ line used in our study marked subpopulations of ocular surface epithelium, which significantly differ from the previously reported K14^CreER^ pattern with more uniform labeling (Richardson, et al., 2017; Amitai-Lange, et al., 2015; Di Girolamo, et al., 2015). Interestingly, a less efficient K14^CreER^ line allowed us to achieve enriched labeling of IFE basal layer away from hair follicles and identify a subpopulation of epithelial cells with relatively higher K14 promoter activity (Sada, et al., 2016; Zhang, et al., 2010). The differences in the K14^CreER^ labeling pattern may also be attributed to the Cre reporter line used in our and previous studies: Rosa-tdTomato shows higher activity than other reporters, including Confetti. Our results suggest that the combination of K14^CreER^ and reporter line may provide effective tools to distinguish different compartments of cornea and conjunctiva.

The SC potential of the central cornea has been under debate. Our lineage tracing showed that K14^CreER+^ and Dlx1^CreER+^ cells in the central corneal compartment are mostly shorter-lived, which might fit the definition of progenitor cells. However, the rare existence of long-lived populations in the central cornea may reflect the SC leakage phenomenon as previously suggested (Lobo, et al., 2016).

After the damage of SCs and their niches, the SC compartment is remodeled depending on the severity of injuries, i.e., localized vs. diffuse and superficial vs. deep, and different SC populations react differently to repair damaged epithelium. We demonstrated that Slc1a3^CreER+^ limbal/peripheral corneal vs K14^CreER+^ central corneal populations indeed respond differently to the limbal deletion, possibly due to the differences in their intrinsic properties or external conditions along the spherical axis of the ocular epithelium. Our K14^CreER^ lineage tracing also showed that conjunctival SCs in the bulbar region, even though they are located closely to the limbus, do not respond to the limbal epithelial damage, indicating distinct responsiveness of corneal and conjunctival SCs to the limbal deletion. In contrast, chemical burn, an experimental model of limbal SC deficiency, triggers invasion of conjunctival SC populations to the cornea without adapting their fates to corneal lineages. This behavior is different from hair follicle SCs transiently recruited to the IFE upon injury, changing their fates to IFE lineage (Ito, et al., 2005). Our genetic tools may provide useful tools to further address cellular and molecular mechanisms of SC plasticity in different SC models and to find potential therapeutic strategy for limbal SC deficiency.

## Experimental Procedures

### Mice

All mouse experiments were performed according to the protocols approved by Animal Care and Use Committee of University of Tsukuba and Kumamoto University. Mice were housed in Laboratory Animal Resource Center, University of Tsukuba, and the Center for Animal Resources and Development in Kumamoto University. For lineage tracing, Slc1a3^CreER^ (C57BL/6J) (Nathans, 2010) (The Jackson Laboratory, no. 012586), Dlx1^CreER^ (C57BL/6J) (Taniguchi, et al., 2011) (The Jackson Laboratory, no. 014551) or K14^CreER^ (a mixed background of CD1 and C57BL/6J) (Vasioukhin, et al., 1999) (a gift from E. Fuchs, Rockefeller University, New York, USA), were crossed with Rosa-tdTomato reporter mice (C57BL6/J) (Madisen, et al., 2010) (The Jackson Laboratory, no. 007905). C57BL/6J wild-type mice were purchased from Charles River Laboratories or Japan SLC, Inc. Adult male and female mice at age of 2-3 months were used for experiments.

### EdU and Tamoxifen treatment

To label LRCs, EdU (50 μg/g body weight, Invitrogen) was injected intraperitoneally twice a day for 1 week, followed by 5 weeks of chase without EdU before euthanization.

For lineage tracing using K14^CreER^, a single dose of Tamoxifen (50 μg/g body weight, Sigma) was injected intraperitoneally at 2-3 months of age. For Slc1a3^CreER^ and Dlx1^CreER^ lines, mice were injected with Tamoxifen (100 μg/g body weight) for 5 consecutive days. Mice were euthanized at the indicated chase periods after the last injection. CreER/Rosa-tdTomato mice without Tamoxifen injections were used to examine the leakiness of Cre.

### Staining of eye sections

Enucleated eyes were fixed in 4% paraformaldehyde (PFA) overnight and snap-frozen in OCT compound (Tissue-Tek, Sakura). For histological analysis, 10-µm sections were air dried and washed, followed by stained with Hematoxylin (Wako, 131-09665) for 20 minutes and Eosin Y (Wako, 058-00062) for 15 seconds. Sections were dehydrated and mounted in Entellan new mounting solution (Merck Millipore, HX73846161). For immunostaining, frozen sections were incubated with blocking solution (2.5% donkey serum and 2.5% goat serum in PBS) for 1 hour at room temperature. Primary antibodies were used at the following dilutions: rabbit anti-K12 (1:100, Abcam, ab185627), rabbit anti-K19 (1:100, Abcam, ab52625). Secondary antibodies (Alexa 488 or 546, Invitrogen) were used at 1:200 dilution. All samples were counterstained with Hoechst (Sigma, B2261) for 10 minutes and mounted. Preparations were analyzed and imaged by Zeiss, Axio Imager.Z2. The brightness and contrast of images were adjusted with equal intensity among different mouse groups by using Adobe Photoshop.

### Whole-mount immunostaining

Eye was cut in half and incubated in EDTA (20 mM)/PBS in an orbital shaker at 37°C for 2 h to separate the epithelium from the mesenchyme as an intact sheet. Epithelial sheets were fixed in 4% PFA overnight at 4°C with gentle shaking. The epithelial sheets were washed, incubated in blocking buffer (1% BSA, 2.5% donkey serum, 2.5% goat serum, 0.8% Triton in PBS) for 3 h at room temperature, and incubated with primary antibodies/blocking buffer overnight at room temperature. Samples were washed 4 times in 0.2% Tween/PBS for 1 h at room temperature, and were incubated overnight with secondary antibodies at 4°C. After washing, samples were counterstained with Hoechst (Sigma, B2261) for 1 h and mounted. Primary antibodies were used at following dilutions: rabbit anti-K12 (1:300, Abcam, ab185627), rabbit anti-K19 (1:300, Abcam, ab52625). Secondary antibodies (Alexa 488, 546 or 647, Invitrogen) were used at 1:200 dilution.

For EdU staining, the epithelial sheets were blocked in blocking buffer for 3 h at room temperature and incubated with the Alexa Flour 488 Click-iT EdU Imaging Kit (Invitrogen) for 1 h at room temperature. Samples were further washed 3 times for 15 min in 0.2% Tween/PBS at room temperature. Samples were co-stained with primary and secondary antibodies as described above. To recover tdTomato fluorescence, samples were incubated 3 times in 0.1 M EDTA for 20 min, followed by 5 min wash in PBS before mounting.

Whole-mount preparations were analyzed and imaged by Zeiss, Axio Imager.Z2 or confocal microscopy (Zeiss LSM710). All confocal data are shown as projected Z-stack images.

### Injury

Limbal deletion and chemical injury was performed according to the previous report (Nasser, et al., 2018). For all injury experiments, Avertin was used for the mouse anaesthesia. Mice were intraperitoneally injected Carprofen (5mg/kg, Rimadyl, Zoetis) and monitored for pain and eye infections during and after injury procedures. The limbus epithelium located above the vessels of the stroma was deleted using an ophthalmic rotating burr (Algerbrush) under Stereo microscope (Zeiss). To verify the complete removal of the epithelium, a drop of 1 mg/mL fluorescein sodium (Sigma, F6377) was applied onto the cornea. For chemical injury, 3 µl of 1 N sodium hydroxide solution was applied to the cornea. Subsequently, the eye was washed with PBS.

### Quantification of microscope images

The length between the corneal/conjunctival boundary and the proximal edge of each clone was measured in the whole-mount samples using ImageJ (Fiji) software. Distribution percentage were calculated by using GraphPad Prism 8 software. All quantifications were independently performed on ≥3 mice. Data are shown as means ± standard error of the mean (S.E.M.).

### Reproducibility

All experiments with or without quantification were independently performed at least three times with different mice and the representative data are shown.

## Acknowledgements

We thank Drs. Keisuke Okabe and Yoshiaki Kubota (Keio University) for teaching us histological analysis of mouse eyes. This work was supported by AMED-PRIME, AMED (JP20gm6110016) and research grants from The Ichiro Kanehara Foundation for the Promotion of Medical Sciences and Medical Care to A.S.

## Author Contributions

A.S., H.Y., and R.I. conceptualized the project. A.S. provided knowledge and techniques of stem cell analysis. R.I. and A.S. performed experiments and analyzed the results. A.S. and H.Y. interpreted the results and supervised the project. R.I., A.S., and H.Y. wrote the manuscript. A.S. acquired funding.

## Declaration of Interests

The authors declare no competing interests.

**Supplementary Figure 1**. Tamoxifen-inducible Cre for cell fate tracking. (A) Immunostaining of wild-type mouse eye on section (left) or whole-mounts (right). K12 and K19 (green) are used as markers of the corneal and conjunctival epithelium, respectively. Hoechst nuclear staining in blue. Scale bars: 500 μm. (B) Schematic representation of Tamoxifen (TM)-inducible CreER system. (C, D) Scheme for long-term lineage tracing. Schedule of TM injection (red arrowheads) and tissue collection (white arrowheads) is shown. (E) K14^CreER^, Dlx1^CreER^ and Slc1a3^CreER^ without Tamoxifen injection. Red, tdTomato. Green, K12 (corneal marker). Blue, Hoechst. Scale bars: 500 μm. (F) Diagram shows the measurement of the length between the peripheral edge of tdTomato+ clone and corneal/conjunctival boundary.

**Supplementary Figure 2**. Examples of lineage tracing of Slc1a3^CreER^ for 1 year. (A, B) Mouse eyes are analyzed by whole-mount immunostaining at 1-year-chase. The yellow dashed line represents corneal/conjunctival boundary. The white dashed line represents dTomato+ cell-enriched area in the fornix conjunctiva. Central corneal areas are shown with higher magnification (right top) and side view of Z-stack confocal image (right bottom). White arrowheads indicate tdTomato+ radial stripes extended from the limbus. Yellow arrowheads indicate tdTomato+ clones expanded laterally within the limbal region. Red, tdTomato. Green, K12 (A, corneal marker). Green, K19 (B, conjunctival marker). Blue, Hoechst. Scale bars: 500 μm (left), 200 μm (right top), 20 μm (right bottom).

**Supplementary Figure 3**. Lineage tracing of Dlx1^CreER^ in the cornea. (A-C) Mouse eyes are analyzed by whole-mount immunostaining at 2-week-, 1-month-, 3-month- and 1-year-chases. Central corneal areas are shown with higher magnification (B) and side view of Z-stack confocal image (C). The yellow dashed line represents corneal/conjunctival boundary (A). Arrowheads indicate tdTomato+ cells (A, B). Red, tdTomato. Green, K12 (B, C, corneal marker). Blue, Hoechst. Scale bars: 500 μm (A), 200 μm (B), 20 μm (C). (D) Number of tdTomato+ clones per a half whole mount sample is quantified at indicated time points. (E) Box plot shows the length of tdTomato+ clone from the corneal/conjunctival boundary at indicated time points of chase. *N* ≥ 3. All tdTomato+ clones in whole-mount samples from a half eye are measured and used for quantification. Error bars show S.E.M.

**Supplementary Figure 4**. Time course of injury experiments. (A) Whole-mount immunostaining after limbal epithelial deletion. 1-day-, 2-week-, and 4-week-post-injury are shown. The yellow dashed line represents corneal/conjunctival boundary. The white dashed line represents dTomato+ cell-enriched area. White arrowheads indicate tdTomato+ radial stripes extended from the limbus. Yellow arrowheads indicate tdTomato+ clones expanded laterally within the limbal region. Red, tdTomato. Green, K19 (conjunctival marker) or K12 (corneal marker). Blue, Hoechst. Scale bars: 500 μm (top), 200 μm (bottom). (B) Whole-mount immunostaining after chemical burn. 2-week-post-injury are shown. White arrows represent movement of conjunctival tdTomato+ clones. Red, tdTomato. Green, K12 (corneal marker) or K19 (conjunctival marker). Blue, Hoechst. Scale bars: 500 μm.

