## Supplementary figures and images for "Defining compartmentalized stem and progenitor populations with distinct cell division frequency in the ocular surface epithelium"

### Supplemental Figure1

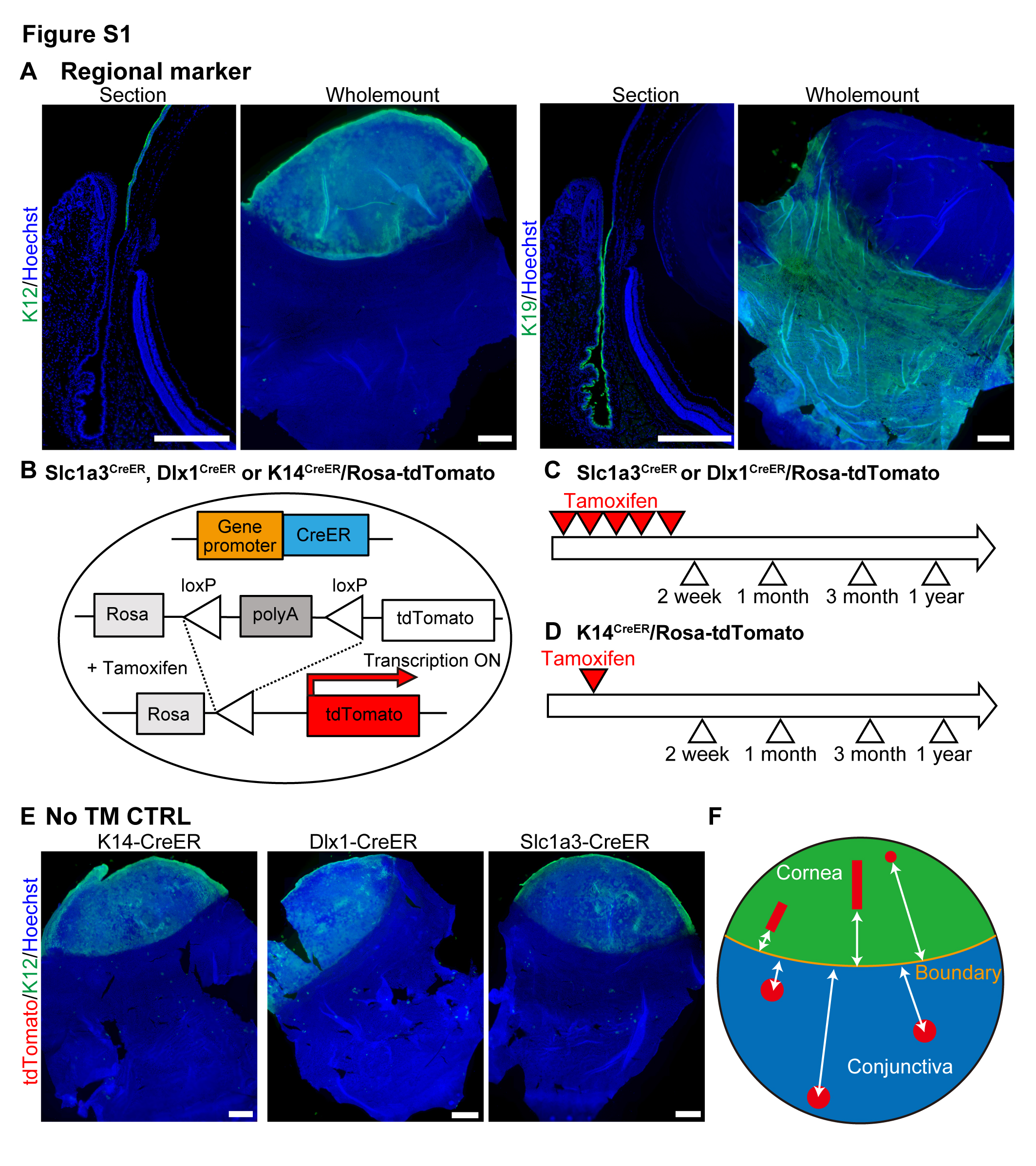

### Supplemental Figure2

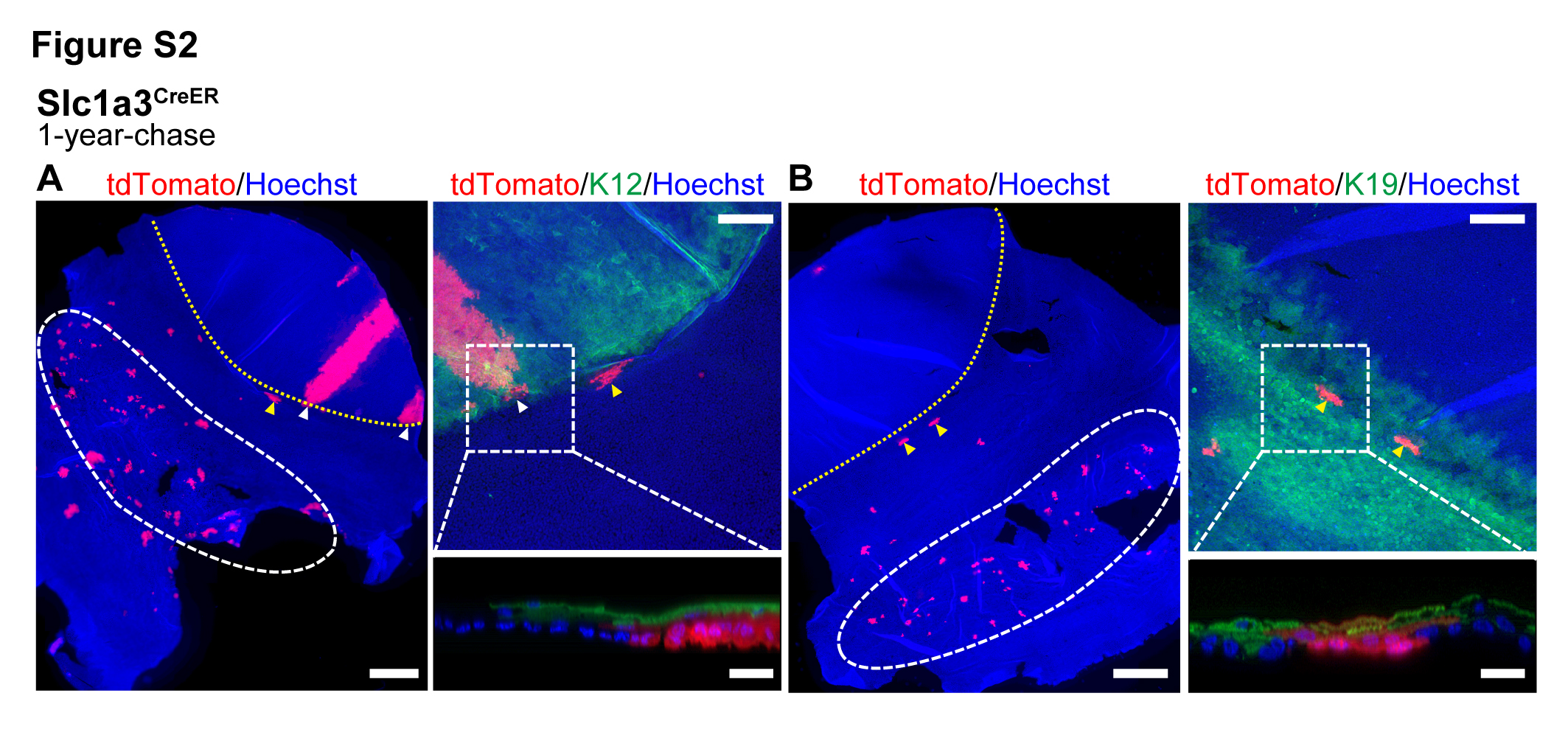

### Supplemental Figure3

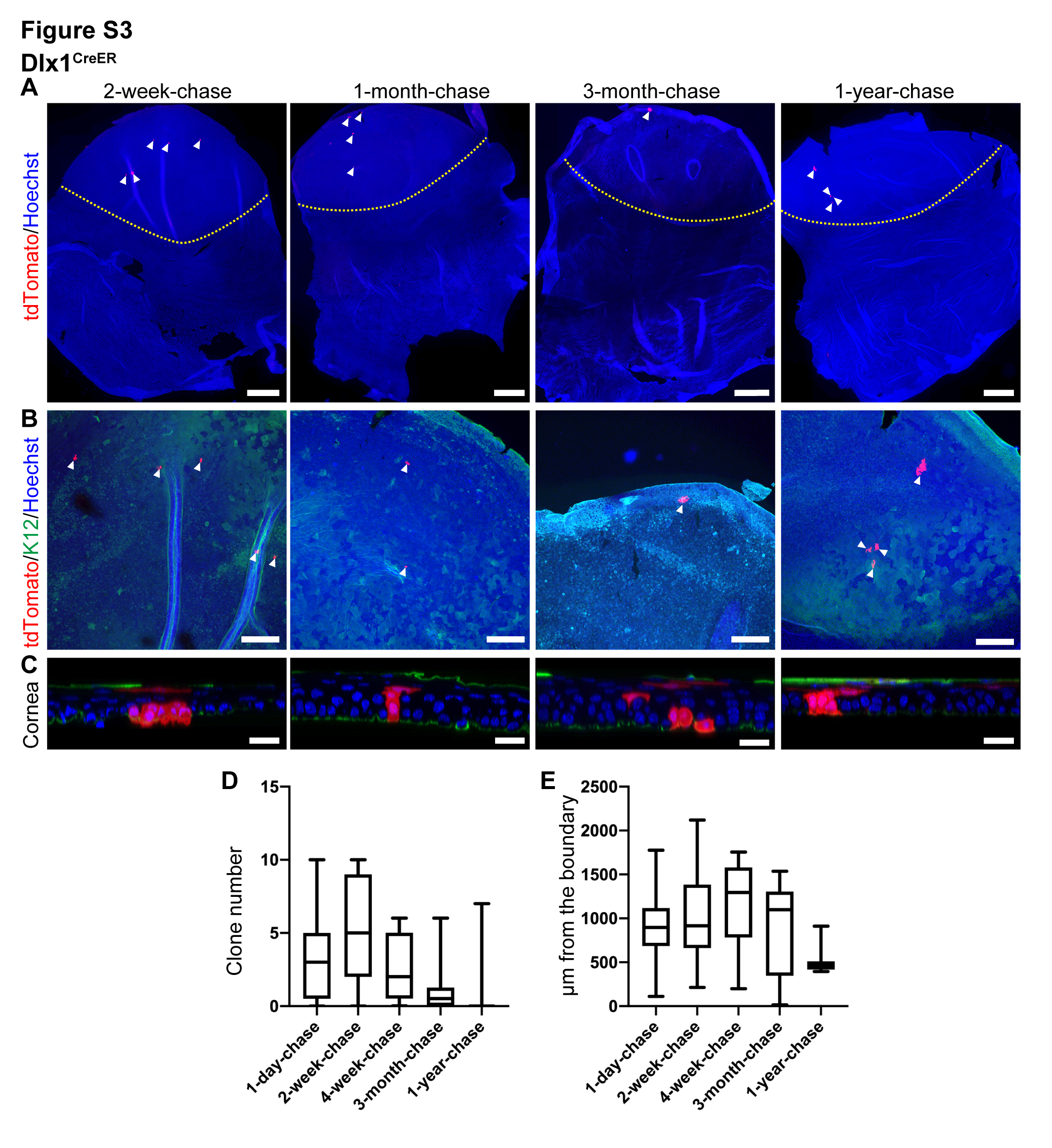

### Supplemental Figure4

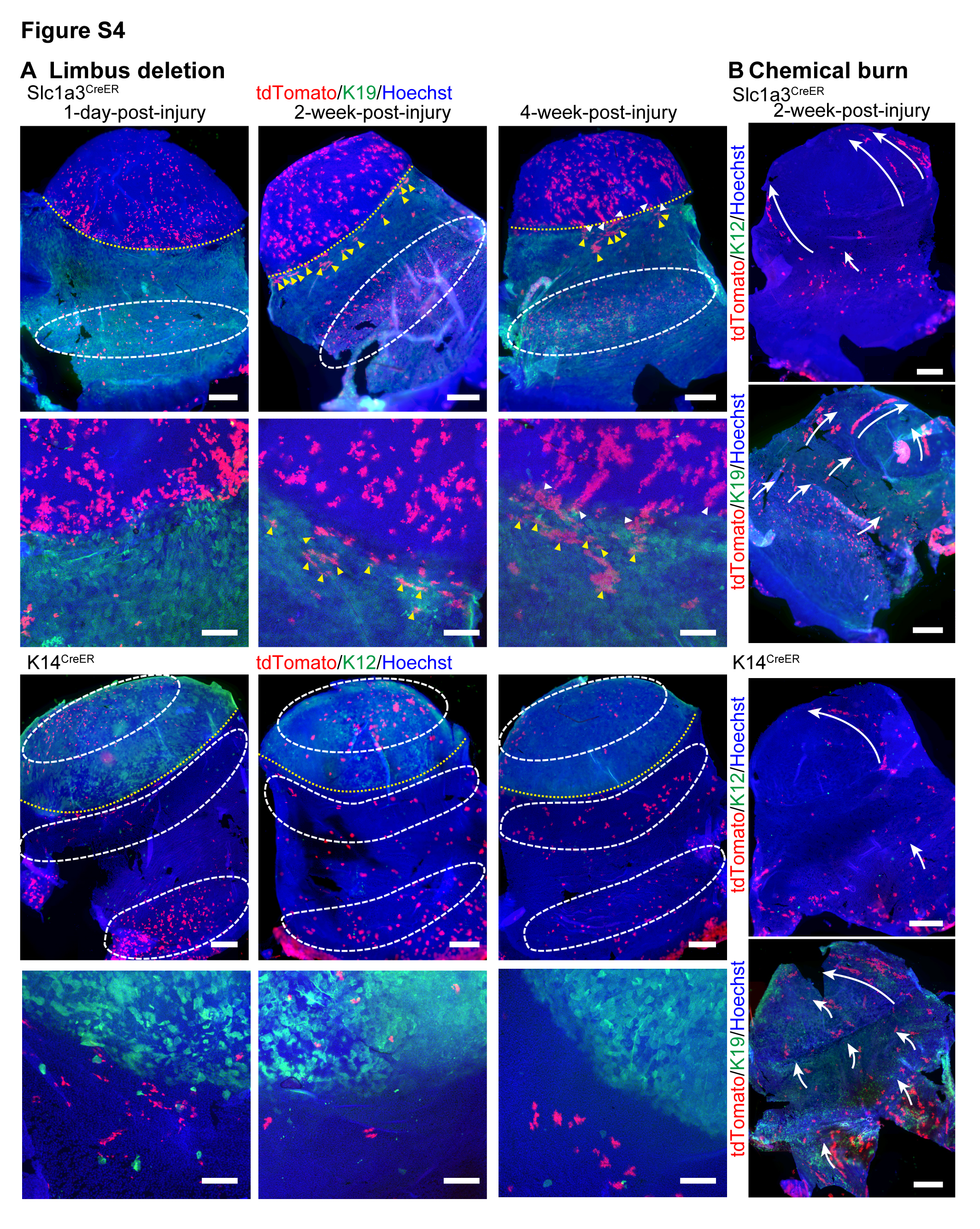
